# Magnetic Bead-Quantum Dot (MB-Qdot) CRISPR Assay for Instrument-Free Viral DNA Detection

**DOI:** 10.1101/2020.04.10.036558

**Authors:** Mengdi Bao, Erik Jensen, Yu Chang, Grant Korensky, Ke Du

## Abstract

We have developed a novel detection system which couples CRISPR-Cas recognition of target sequences, Cas mediated nucleic acid probe cleavage, and quantum dots as highly sensitive reporter molecules for instrument-free detection of viral nucleic acid targets. After target recognition and Cas mediated cleavage of biotinylated ssDNA probe molecules, the probe molecules are bound to magnetic particles. A complimentary ssDNA oligonucleotide quantum dot conjugate is then added, which only hybridizes to un-cleaved probes on the magnetic beads. After separation of hybridized from unhybridized quantum dot conjugates by magnetic sequestration, the signal is measured fluorometrically to provide a signal proportional to the cleaved probes and therefore the amount of target nucleic acid. To demonstrate the power of this assay, a 250 bp DNA target sequence matching a portion of the African swine fever virus (ASFV) genome is used and a detection limit of ~0.5 nM is achieved without target amplification using a simple portable ultraviolet flashlight. The positive samples are readily confirmed by visual inspection, completely avoiding the need for complicated devices and instruments. This work establishes the feasibility of a simple, instrument free assay for rapid nucleic acid screening in both hospitals and point-of-care settings.

## Introduction

Intensified infectious disease outbreaks have occurred more frequently in the past few decades due to the increased human population and global travel, and have included viruses as SARS^1,2^, Ebola^3,4^, Zika^5,6^, MERS^7,8^, and the novel coronavirus (SARS-CoV-2)^9,10^. The ongoing COVID-19 crisis has evolved as a global pandemic since it was first reported in December 2019. Due to the shortages of testing kits, vaccines, and treatments, the number of cases is still rapidly increasing in many countries. Since the development of vaccines and treatments for novel viruses can take as long as 18 months, the most efficient immediate approach to slow down and contain epidemics is to provide rapid and widespread testing, followed by the isolation of positive cases and their close contacts. The current standard tests for viral infection use highly sensitive and specific real-time polymerase chain reaction (RT-PCR)^11,12^. Although these tests are highly sensitive and accurate, they require a laboratory setting for use, bulky instruments, sophisticated operations, and they are therefore not field deployable. These issues present a significant bottleneck to rapid and widespread testing, hindering our ability to respond to emerging pandemics. Recently, rapid POC systems using isothermal amplification have been demonstrated for viral pathogen detection^13^, however they have limited throughput and poor sensitivity. Given the limitations of existing viral pathogen testing methods, there is a significant need for simpler testing methodologies that do not require complex instrumentation, and that can be performed easily in the field.

An ideal POC device that can combat modern era pandemic crises should be simple, inexpensive, accurate, and must be easily operated and understood by people without special training. A recently discovered technology called clustered regularly interspaced short palindromic repeats (CRISPR) in conjunction with CRISPR-associated proteins (Cas) has shown great promise as an alternative to RT-PCR detection^14,15^. This method is significantly simpler than RT-PCR since it does not require complex instrumentation to perform. Certain Cas nucleases such as Cas12a and Cas13a can be activated to indiscriminately cleave single-stranded DNA or RNA substrates once they identify and bind with the target DNA or RNA^16,17^. The activated Cas enzymes cleave a nucleic acid probe, releasing a fluorophore from a quencher molecule, thereby generating signal. CRISPR-Cas systems have also been linked with isothermal amplification methods to achieve highly sensitive and specific detection of pathogens^18,19^. However, most CRISPR-Cas detection systems utilize reporter probes with organic dyes that require external instruments and possess a high fluorescence background, which limits overall sensitivity^20^.

In this study, we develop a novel probe system for CRISPR-Cas nucleic acid assays which use quantum dot as a reporter. Compared to conventional organic fluorescence dyes, Qdots have unique electronic and optical properties such as stable signal intensity, narrow emission spectrum, continuous broad excitation spectrum, and high quantum yields, making them an ideal fluorescence indicator for in vitro diagnostics^21,22^. After target recognition and Cas mediated cleavage of biotinylated ssDNA probe molecules, the probe molecules are bound to streptavidin-coated magnetic particles. A complimentary ssDNA oligonucleotide quantum dot conjugate is then added, which only hybridizes to un-cleaved probes on the magnetic beads. After separation of hybridized from unhybridized quantum dot conjugates by magnetic sequestration, the signal is measured fluorometrically to provide a signal proportional to the cleaved probes and therefore the amount of target nucleic acid. Our use of magnetic particle sequestration effectively reduces background signal, enabling use highly sensitive quantum dots as reporters. As a proof of principle for viral pathogen detection, a double-stranded DNA segment matching a portion of the African Swine Fever Virus (ASFV) genome was selected as the detection target. Without target amplification, we achieved a detection limit of 0.5 nM by using a small and inexpensive UV flashlight as the excitation source. This simple and instrument-free protocol pave the way for next-generation POC and field deployable viral pathogen detection applications.

## Results

**Fig. 1a** shows the workflow of the Qdots-CRISPR based fluorometric nucleic acid detection. Oligonucleotide DNA linker probes with a biotin label are mixed with a Cas12a-crRNA complex. With the presence of a double-stranded target DNA (part of the ASFV genome), the Cas12a-crRNA complex is activated and indiscriminately cleaves the single-stranded linker probes in the assay. After the reaction, streptavidin-coated magnetic beads are added which bind both cleaved and uncleaved linker probes. A complimentary oligonucleotide DNA reporter probe which is conjugated to quantum dots is then added. The reporter probe only binds to uncleaved linker probes on the magnetic beads. An external magnet is then applied to isolate the magnetic particles, leaving the unbound reporter probes in the eluate for fluorescence analysis. Since increasing the cleavage of the linker probe increases the amount of reporter probe remaining in the eluate, the fluorescence signal is proportional to the amount of starting target in the sample. Transmission electron microscopy (TEM) images of the streptavidin-coated quantum dots (mean diameter of 17.5 nm), streptavidin-coated magnetic beads (mean diameter of 1 μm) and hybridization of the Qdot reporter probe onto linker probes bound to the magnetic beads are presented in **Fig. 1b**. Numerous Qdots are immobilized on the magnetic bead by the complementary hybridization of reporter probe and linker probe. As shown in **Fig. 1c**, Qdots exhibit higher fluorescence intensity than conventional organic fluorescein amidite (FAM) (excitation: 488 nm), making them ideal for instrument-free visual detection. **Fig. 1d** further shows the workflow of our detection scheme: cleaved and uncleaved linker probes are immobilized on the magnetic beads, followed by the hybridization with Qdot reporter probe. Magnetic isolation separates unhybridized reporter probes, leaving probes in the eluate that are proportional to the starting target nucleic acid. Fluorescence measurement then can be performed with the illumination of an inexpensive and portable flashlight.

**Figure 1.**
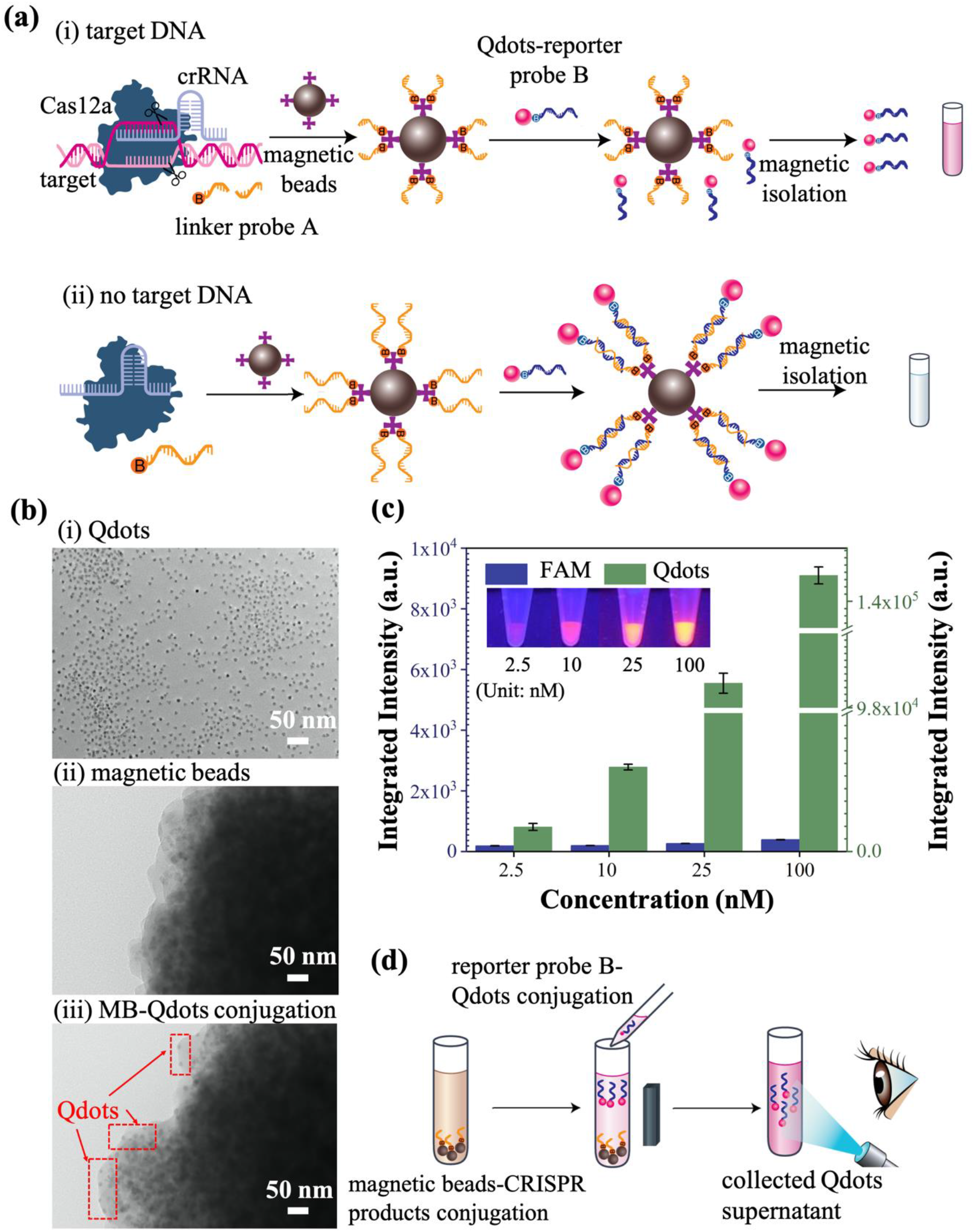
(a) Scheme for the Qdots-CRISPR based nucleic acid detection strategy. (b) TEM images: (i) streptavidin-coated Qdots; (ii) streptavidin-coated magnetic beads; and (iii) MB-Qdot conjugation. The MB-Qdot conjugation includes magnetic beads, DNA probes, and Qdots. Scale bars are 50 nm. (c) Integrated fluorescence signals of FAM organic dye versus Qdots (excitation: 480 nm). Inset shows the photographs of Qdots with a concentration of 2.5 nM, 10 nM, 25 nM, and 100 nM, excited by a portable UV flashlight (wavelength: 395 nm). Error bars represent standard deviation of the mean. (d) Schematic of the instrument-free detection of using Qdots.

A custom designed fluorometer was used to confirm the successful formation of conjugates. As shown in **Fig. 2a**, streptavidin-coated Qdots (2.6 nM) show a fluorescence intensity of ~24 counts (Qdot: green). MB-Qdots conjugation comprised of magnetic beads (100 μg), linker probes A (25 pmoles), reporter probes B (12.5 pmoles), and Qdots (156 fmoles) was mixed in the solution of 60 μL and pulled down by a magnet. Since the input Qdots were saturated by the other three components, no fluorescence signal was detected in the supernatant (Conjugation: red). To confirm the MB-Qdot conjugation was made by the hybridization of two DNA strands, we mixed magnetic beads, reporter probes, and Qdots without the input of linker probes. After magnetic isolation, a high fluorescence intensity (~14 counts) was detected, indicating that the MB-Qdot synthesis is dominated by DNA hybridization rather than non-specific absorption (Control: blue). To further confirm the MB-Qdots conjugation, the beads were washed 5 times followed by DNase I (EN0521, ThermoFisher Scientific) treatment to degrade the hybridized DNA strands. As shown in **Fig. 2b**, a distinct fluorescence peak was detected for the conjugation sample (red), while no signal was observed in the control sample (streptavidin-coated magnetic beads). The integrated signals of the fluorescence spectra (550-700 nm) were plotted (inset of Fig.2b), where the conjugation sample shows ~2,000 counts, and is approximately 13 times higher than the control sample. In addition, for the conjugation sample, the bright pink fluoresence signal is easily observed by using a flashlight.

**Figure 2.**
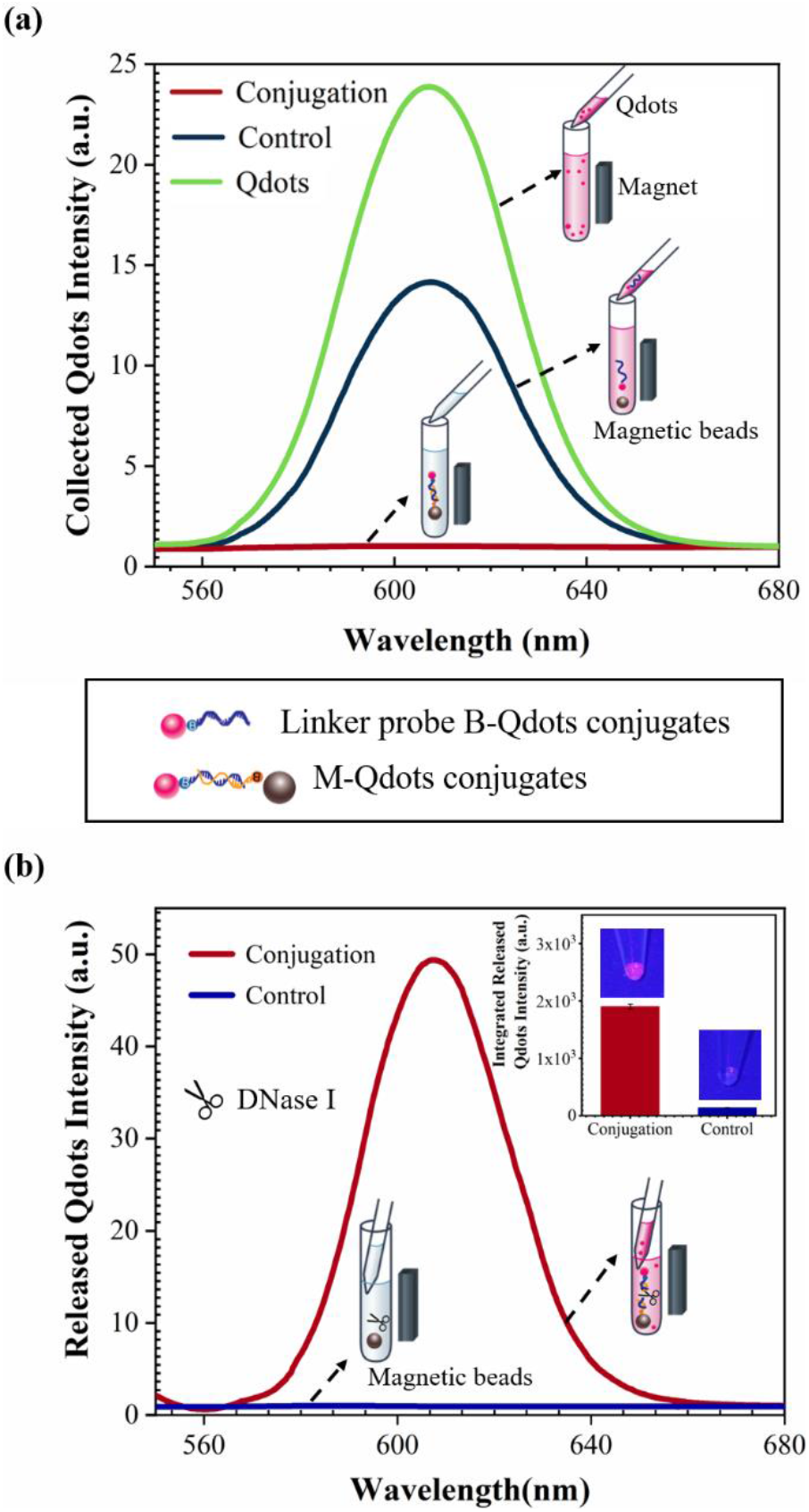
Validation of the MB-Qdot conjugation formation. (a) Emission curves of collected supernatants. The conjugation sample consists of magnetic beads, DNA probes, and Qdots. The control sample consists of magnetic beads, Qdots, and reporter probes. The Qdots sample is the streptavidin-coated Qdots (2.6 nM). (b) Fluorescence signals of released Qdots by DNase treatment versus streptavidin-coated magnetic beads. Inset shows the integrated fluorescence signals and photographs of the suspensions under UV illumination. Error bars are standard deviation of the mean.

We then optimized the MB-Qdot conjugation assay to decrease the fluorescence background, which is essential for the CRISPR Cas12a-based detection strategy. Linker probes (100 pmoles) with 3’ FAM modification (/5BioTinTEG/TTATTCTTATTGTGTGAACTGCTCCTTCTTGACTCCACC/36-FAM/) were introduced to bind with the magnetic beads at weights of 0, 25, 100, 400, and 600 μg at room temperature for 15 min. After incubation, the magnetic beads were isolated, and the fluorescence intensity of the unbounded ssDNA was measured using a commercial spectrofluorometer (JASCO FP-8500, USA). As shown in **Fig. 3a**, the integrated fluorescence intensity (500 to 700 nm) dramatically drops with the increase of the magnetic beads input from 0 μg to 400 μg. Adding more magnetic beads does not decrease the fluorescence intensity, indicating that the saturated ratio of magnetic bead to linker probe A is ~4 μg: 1 pmoles. Next, we optimized the ratio of reporter probes B to Qdots, which is the key to eliminate the fluorescence background. One pmoles of Qdots were selected and the load of reporter probe B was varied from 0 to 80 pmoles. As shown in **Fig. 3b**, a ratio of 80:1 between reporter probe B and Qdots shows the lowest background and is used for nucleic acid detection. To determine the optimized hybridization condition of linker probe A and reporter probe B, Qdots with a load of 0.5 pmoles were pre-mixed with 40 pmoles of reporter probe B (ratio of 1: 80), and then allowed to hybridize with magnetic bead-linker probe A conjugation (ratio of 4 μg: 1 pmoles). The retrieved Qdots were then measured by a custom designed fluorometer. The fluorescence intensity of the supernatant drops significantly by increasing the load of linker probe A from 20 to 50 pmoles (**Fig. 3c**). However, the intensity only shows a slight change when the load is further increased to 80 and 100 pmoles. Thus, we concluded that the saturated ratio of linker probe A to reporter probe B is ~2:1.

**Figure 3.**
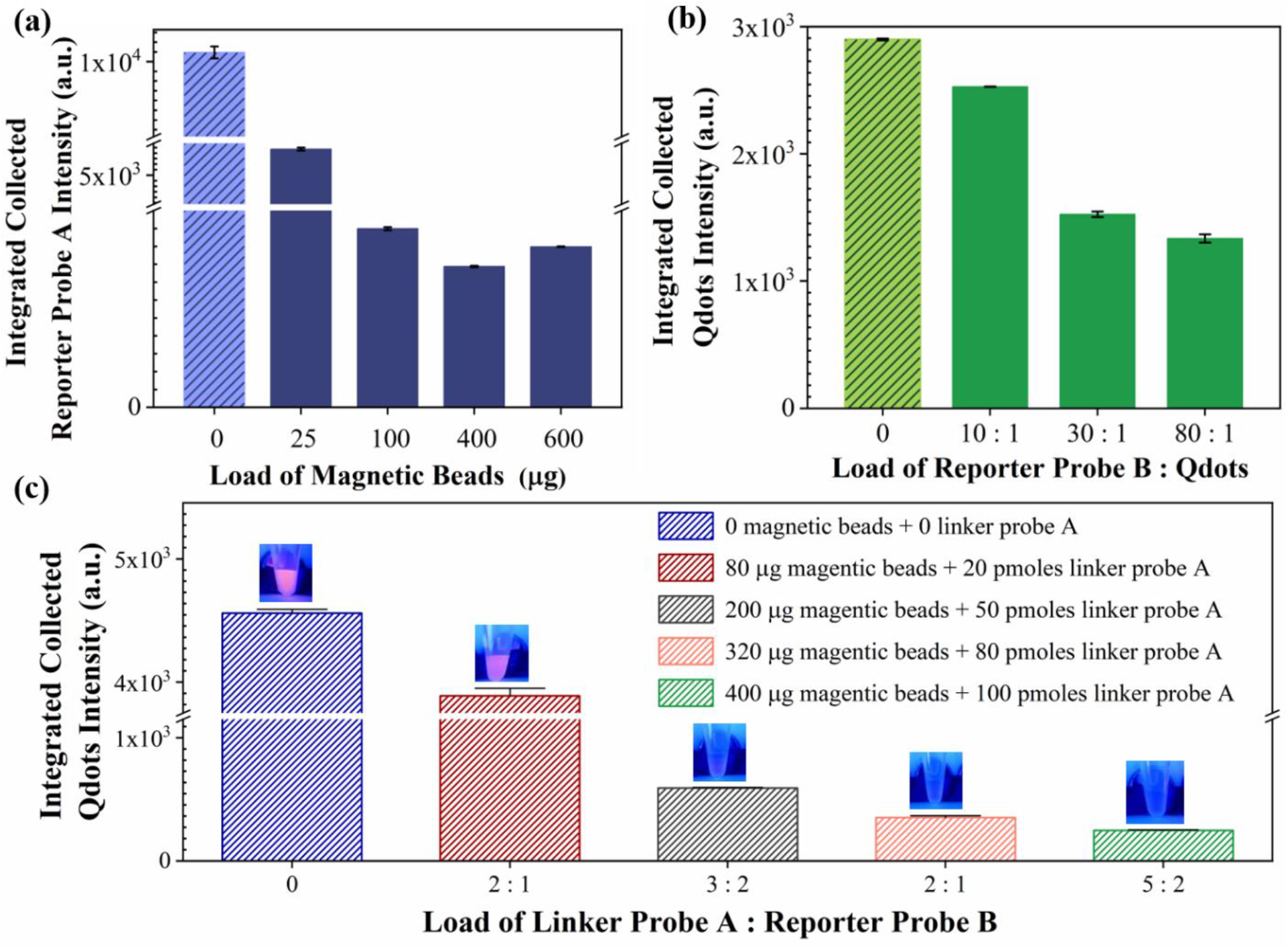
Optimization of the MB-Qdot conjugation. (a) Integrated fluorescence signal of unbound linker probe A versus the input load of magnetic beads (0-600 μg). (b) Integrated fluorescence signal of unbound Qdots versus the input load of reporter probe B (0-80 pmoles). (c) Integrated fluorescence signals of collected Qdots supernatant versus the input load of linker probe A (0-100 pmoles). Inset shows the photographs of Qdots under UV illumination. Error bars represent standard deviation of the mean.

With the optimized MB-Qdot conjugation assay in hand, we then applied this protocol for instrument-free CRISPR detection. We added target DNA into the tube containing 50 nM of Cas12a, 62.5 nM of crRNA, and 1.25 μM of linker probe A with a total volume of 20 μL. As presented in **Fig. 4a**, under flashlight illumination, a distinct pink color is observed with a target concentration higher than 0.5 nM. On the other hand, the collected supernatants do not show any fluorescence signals without target input or with a mismatched target sequence. These results were then validated by using the custom designed fluorometer and are presented in **Fig. 4b**. The intensities of matched target DNA samples with concentrations of 0.5 nM, 0.75 nM, 1 nM, 2.5 nM, 10 nM, and 50 nM are significantly higher than the mismatched or blank samples. These results were further confirmed by TEM imaging. As shown in **Fig. 4c**, for blank and mismatched samples, Qdots with an average diameter of 17.5 nm were linked on the magnetic bead via linker probe A and reporter probe B hybridization. On the other hand, for samples containing target DNA, Qdots are not observed in the TEM image, indicating that the linker probes were denatured and left the Qdots suspended in the solution for detection.

**Figure 4.**
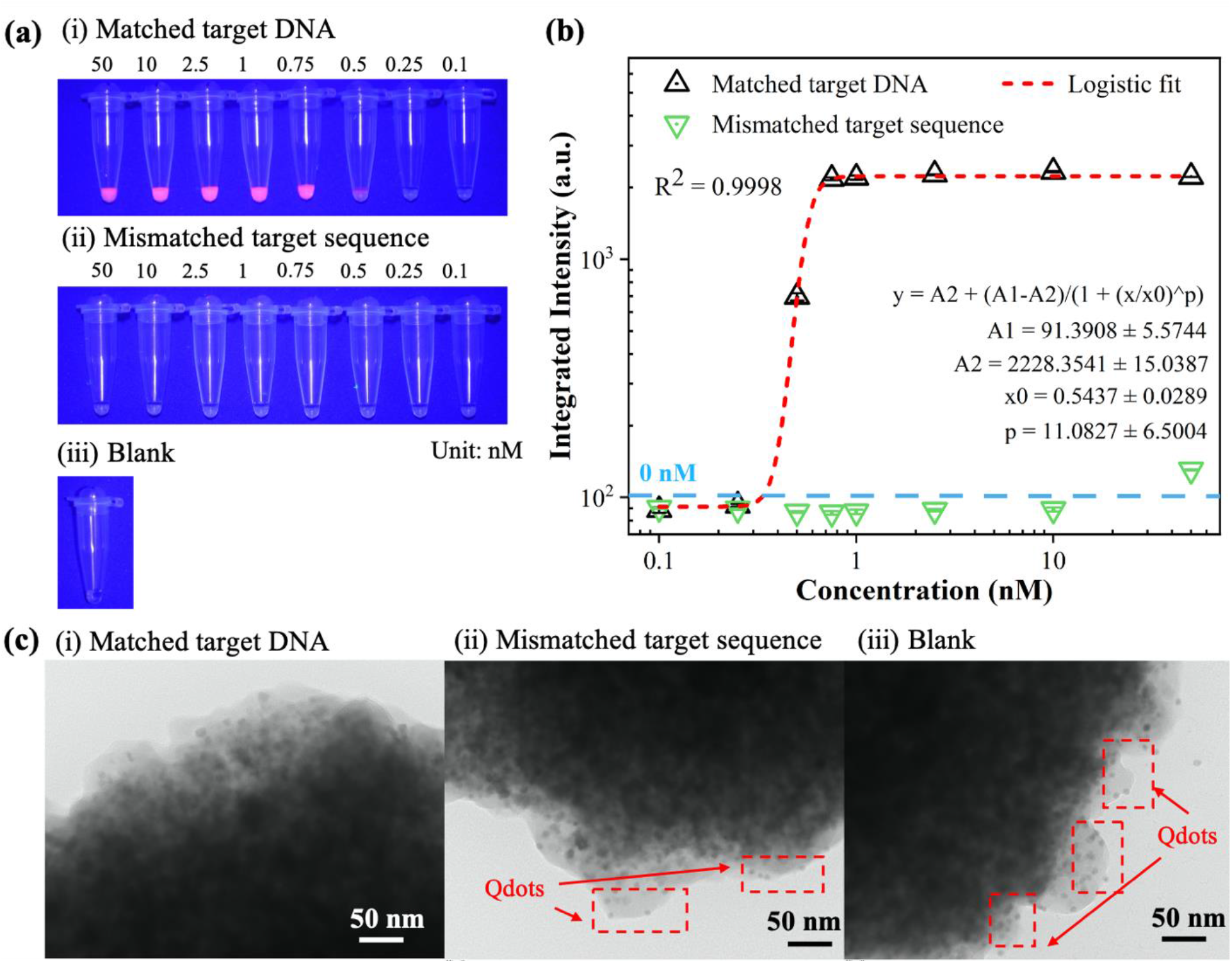
Qdots-CRISPR based nucleic acid detection. (a) Fluorescence images of samples containing (i) matched target DNA (positive); (ii) mismatched target sequence (negative); and (iii) blank (no target input). The samples were illuminated by a UV flashlight (wavelength: 395 nm). (b) Integrated fluorescence signal (550-700 nm) of (i) matched target DNA (black triangle); (ii) mismatched target sequence (green triangle); and (iii) blank sample (0 nM). The red dot line shows the logistic fit of the integrated fluorescence intensity versus DNA concentration. (c) TEM images of the magnetic beads of mixed with (i) matched target DNA; (ii) mismatched target sequence; and (iii) blank (no target input).

## Discussion

The work presented here leverages a CRISPR Cas12a assay and uses Qdots as an indicator. Unlike conventional organic dyes such as FAM, Qdots are brighter due to their high quantum yield and show much better photostability. Coupled with CRISPR Cas12a assay, the reaction is sensitive, rapid, and can be performed in isothermal conditions. Unlike methods such as PCR^23,24^, electrochemical sensing^25,26^, and surface-enhanced Raman spectroscopy^27,28^, our assay does not require time-consuming sample processing, bulky sensing instruments, or complicated expert operation and data interpretation. These unique advantages enable us to demonstrate a simple and instrument-free diagnosis assay, ideal for rapid on-field applications.

Using our Qdot-CRISPR based DNA detection strategy, we report a very low detection background compared to conventional CRISPR-Cas systems that use fluorophore-quencher substrate as reporters. The cleaved ssDNA probes linked to the Qdots are detected by isolating the magnetic beads, thus avoiding the high background issues from the uncleaved fluorophore-quencher probes in the assay. This is essential to enhance the overall detection sensitivity of CRISPR-Cas systems, and is especially useful for instrument-free detection in resource-limited environments^29,30^.

Currently, our approach can achieve a detection limit of ~0.5 nM without any target amplification. To improve the sensitivity, our assay can be easily integrated with established isothermal nucleic acids amplification methods such as recombinase polymerase amplification (RPA)^31,15^ or Loop-mediated isothermal amplification (LAMP)^32,33^ to extend the detection limit to attomolar level. The amplification reagents can be mixed with CRISPR Cas12a protein and crRNA as a “one-pot” assay.

Furthermore, our assay is poised to be developed for high-throughput multiplexing detection purposes. Because CRISPR Cas12a assay is an indiscriminate DNase activity^14,34^, linker probe A and reporter probe B can be used as universal reporters for the detection of various DNA targets by only changing the crRNA sequence to match the target sequence. In addition, Cas13a protein demonstrates a similar indiscriminate RNase activity as Cas12a^35,16^. Thus, our assay presented here should be able to detect both DNA and RNA targets for multiplexing detection.

## Materials and Methods

### CRISPR Cas12a-based linker probe A cleavage

The total volume of CRISPR Cas12a-based cleavage assay was 20 μL including LbCas12a protein (New England BioLabs, Inc.), crRNA, linker probe A, and binding buffer. Briefly, 50 nM of LbCas12a was assembled with 62.5 nM of crRNA at room temperature for 10 min. Afterward, 1.25 μM of linker probe A, 1× Binding buffer, 14.5 μL of Nuclease Free Water (IDT, Inc.), and target DNA (concentration ranging from 0.1 nM to 50 nM) were mixed with the LbCas12a-crRNA complex to initiate the reaction. The mixture was incubated in a water bath at 37 °C for 120 min to ensure optimized cleavage activity. All the nucleic acids used in the assay were purchased from IDT, Inc., and the sequences are given in **Table.1**.

**Table 1.**
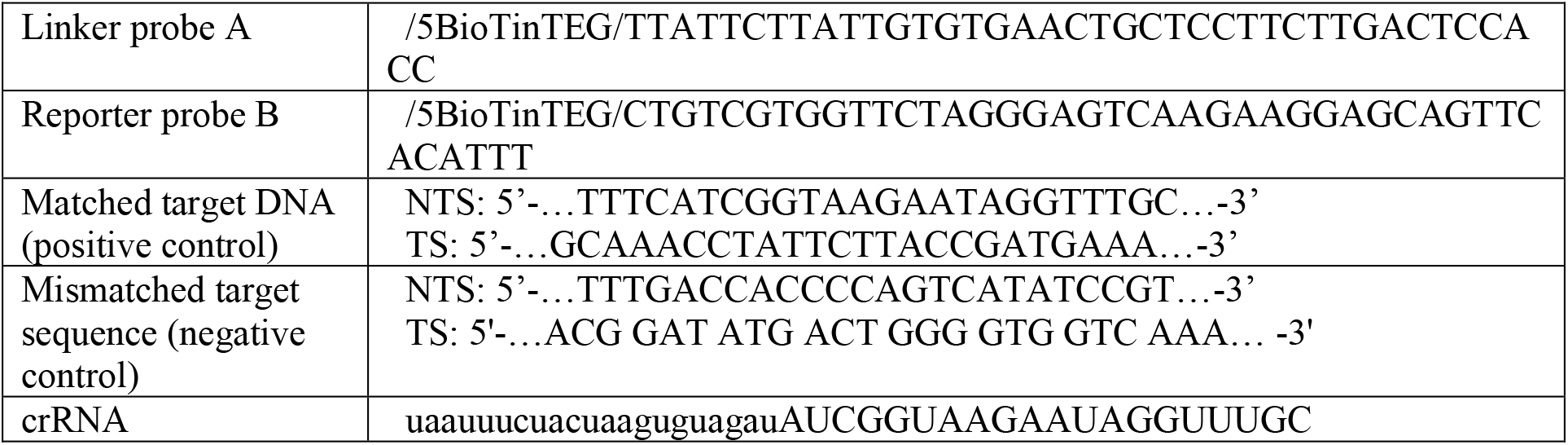
Sequences of DNA and crRNA used in this study.

### Linker probe A isolation via magnetic beads

Firstly, 10 μL of magnetic beads (Dynabeads™ MyOne™ Stretpavidin C1, ThermoFisher Scientific) were washed with PBS buffer (Gibco TM, PH 7.4) for three times and resuspended in 10 μL of the PBS buffer. Next, the washed magnetic beads were transferred to a new 0.2 mL PCR tube, and the supernatant was discarded by isolating the magnetic beads with a magnetic rack (NEBNext® Magnetic Separation Rack, New England BioLabs, Inc.). The collateral cleaved products by CRISPR were then added to the magnetic beads at room temperature for 15 min. After incubation, the magnetic bead-linker probe A conjugation was washed and resuspended in 20 μL PBS buffer for later use.

### Reporter probe B conjugation with quantum dots (Qdots)

Qdots (Qdot™ 605 ITK™ Streptavidin Conjugate Kit, ThermoFisher Scientific) with a concentration of 15.6 nM were mixed with 1.25 μM of reporter probe B (IDT, Inc.) to reach a 10 μL reaction mixture. To avoid photo-bleaching issues, the mixture was incubated in a dark environment at room temperature for 15 min.

### Hybridization of linker probe A and reporter probe B

The supernatant of the magnetic bead-linker probe A conjugation was first evacuated, followed by mixing with the reporter probe B-Qdot conjugation. The mixture was placed on a tube rotator (BS-RTMNI-1, STELLAR SCIENTIFIC) and allowed to react at room temperature under mild shaking for 90 min. After that, the supernatant was collected and transferred to a new PCR tube.

### Naked-eye detection and fluorescence quantification

To identify the presence of target DNA, collected Qdots were illuminated by using a portable flashlight (wavelength: 395 nm). To validate the instrument-free detection, the fluorescence intensity of the Qdots was measured by a custom designed fluorometer. As reported before^34^, the sample was added in a disposable cartridge and excited by a continuous wave semiconductor laser (wavelength: 488 nm). The collected fluorescence signal was analyzed by a portable spectrometer (USB 2000+, Ocean Optics).

### TEM characterization

Transmission electron microscopy (TEM, JEOL 2010) with a point resolution of 0.23 nm was used to characterize the samples. The supernatant containing Qdots was deposited onto a carbon-coated copper grid and left to evaporate. The TEM images were then captured through an AMT XR81 bottom mount camera with a 60 kV electron microscope.

## AUTHOR INFORMATION

### Author Contributions

M. Bao, E. Jensen, and K. Du designed the experiments. M. Bao, Y. Chang, and G. Korensky conducted the experiments. M. Bao and K. Du wrote the manuscript. All authors commented the manuscript.

## ACKNOWLEDGMENT

The authors would like to thank Wenrong He (RIT) for the schematic design.

---

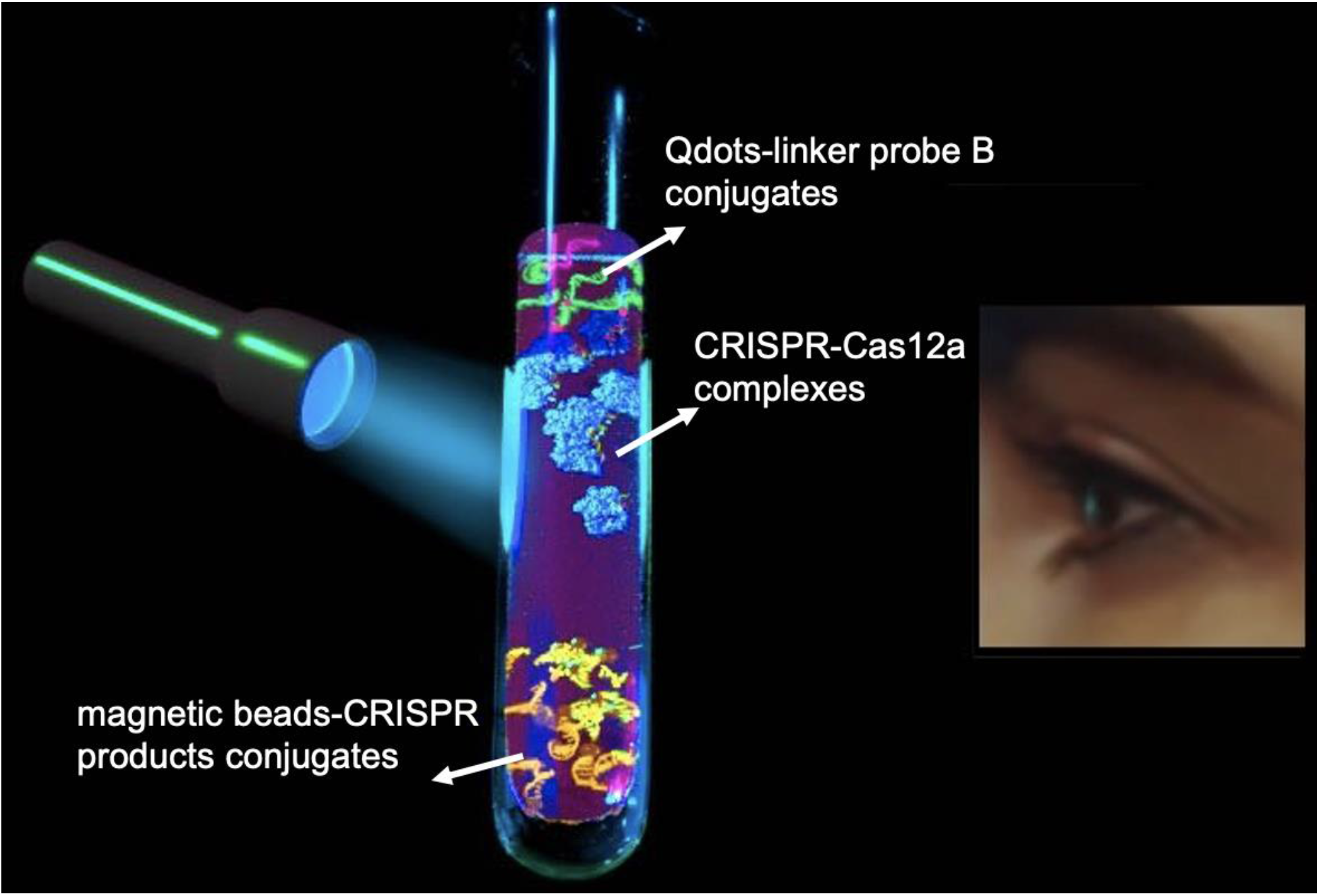
TOC only.

